# Spatio-temporal visualisation and manipulation of endogenous Yap1 in medaka

**DOI:** 10.64898/2026.09.15.751690

**Authors:** Ali Seleit, Yoichi Asaoka, Simon Knoblich, Luc Lederer, Chiharu Oozono, Atsunori Oga, Akihiro Oiso, Makoto Furutani-Seiki, Alexander Aulehla

## Abstract

A key requirement to address fundamental questions in developmental biology is the ability to visualise and manipulate endogenous signalling proteins *in vivo*. Here we generate such an enabling experimental tool for the highly conserved Hippo pathway and its transcriptional effector Yap1, which converts mechanical and chemical cues into transcriptional output during embryogenesis and regeneration. We generate and characterise a Yap1-mGreenLantern (Yap1-mGL) endogenous CRISPR/Cas9-mediated knock-in line that enables 4D analysis of Yap1 dynamics in medaka. We resolve Yap1- mGL expression and nuclear/cytoplasmic localisation in real time, revealing the onset of reporter fluorescence before gastrulation, tissue and cell-type specific Yap1-mGL localisation during somitogenesis and injury-induced Yap1 upregulation during larval spinal cord regeneration. Using the nanobody-based degron system, we degrade the Yap1 protein *in vivo,* recapitulating the genetic loss-of-function mutant phenotype. Combining endogenous tags with nanobody-based degrons offers a powerful approach to visualise and acutely perturb endogenous protein function *in vivo*.

## Introduction

Yap1, a nuclear executor of the Hippo pathway, transduces mechanical and chemical cues into its target gene transcription during vertebrate embryogenesis and regeneration (Dupont et al., 2011; Moya and Halder, 2019; Panciera et al., 2017). Critically, most of its activity is set post-translationally by nuclear (active) *vs.* cytoplasmic (inactive) localisation. Thus, where Yap1 is expressed and where it is active can diverge (Nishioka et al., 2009; Zhao et al., 2007). Current tools to monitor Yap activity largely rely on TEAD-based transcriptional reporters or antibody staining (Astone et al., 2018; Camacho-Macorra et al., 2024; Miesfeld and Link, 2014). To directly monitor Yap protein and subcellular localisation dynamics during embryogenesis and regeneration, our goal was to generate a fluorescently tagged Yap1 protein by genetic engineering at the endogenous locus (Gu et al., 2022). This would serve both the detailed quantification in *time* and *space* in addition to opening possibilities for functional interrogation by nanobody mediated protein degradation strategies (Caussinus et al., 2012; Yamaguchi et al., 2019).

We applied our previously reported CRISPR/Cas9 knock-in protocol (Seleit et al., 2021) to generate a Yap1-mGreenLantern (Yap1-mGL) endogenous fusion protein line. The validation by whole genome sequencing (WGS) confirmed a scarless single-copy in-frame integration at the 3′ end of the medaka *yap1* coding sequence. 4D live imaging during early embryonic development reveals that the onset of detectable Yap1 fusion protein fluorescence signal directly precedes gastrulation in medaka. At somitogenesis stages, we quantify cellular nuclear-to-cytoplasmic (N/C) ratio of Yap1 across the intact tail, enabling a tissue-wide readout of endogenous Yap1 localisation *in vivo.* This revealed that Yap1 is expressed widely in the developing tail but that a high N/C Yap1 ratio occurs primarily in notochordal unipotent progenitors and the epidermis, cell-types that have been shown to be under mechanical tension (Kimelman et al., 2017; Lim et al., 2017; Seleit et al., 2020). Beyond embryogenesis, we report that injury-induced upregulation of Yap1 accompanies wound closure and restoration of tissue continuity following spinal cord transection in medaka larvae, mirroring results described in adult zebrafish (Klatt Shaw et al., 2021; Mokalled et al., 2016), and in contrast to the limited regenerative capacity reported in adult medaka (Aoki et al., 2025). Lastly, we combine the nanobody-based degron system used in zebrafish (Yamaguchi et al., 2019) with our homozygous endogenous Yap1-mGL KI to acutely degrade the Yap1 protein *in vivo,* phenocopying the *hirame* (*yap1*) loss-of-function genetic mutant (Porazinski et al., 2015). Taken together, this work provides a framework in which endogenous protein dynamics and protein function are interrogated through an endogenously tagged fusion protein. Applied to the growing set of knock-in lines in vertebrate models, this approach opens the way to a spatially and temporally resolved understanding of signalling protein function during embryogenesis and in post-embryonic stages.

## Results and discussion

### 4D imaging of endogenous Yap1 dynamics during embryonic development

To generate an endogenously tagged *yap1* in medaka we used our previously described CRISPR/Cas9-mediated knock-in protocol (Seleit et al., 2021) (Supplementary Figure 1A). We generated a *yap1-mGL* knock-in (KI) line by targeting the C-terminus of *yap1* with the sequence coding for the mGreenLantern fluorescent protein (Campbell et al., 2020). Whole-genome sequencing on F2 embryos confirmed scarless single-copy integration of mGL at the *yap1* locus, with no other integration site detected genome-wide (Supplementary Figure 1B-C). Homozygous *yap1-mGL* embryos and larvae do not show any observable phenotypic abnormalities and adults are viable and fertile, indicating that the tag does not disrupt Yap1 function. To assess Yap1-mGL expression dynamics during early medaka development, we performed 4D time-lapse confocal imaging. We detect no maternal Yap1-mGL signal in early medaka embryos. In mouse, maternal Yap1 is provided in the oocyte and is required for activation of the early zygotic genome (Yu et al., 2016), whereas in zebrafish maternal *yap1* transcript is deposited but maternal-zygotic *yap1* mutants are viable, likely reflecting compensation by *taz* (Grimm et al., 2019; Vázquez-Marín et al., 2019). Medaka, by contrast, lacks a functional *taz* but carries the paralogue *yap1b*, which acts as the primary maternal *yap* contribution (Vázquez-Marín et al., 2019). Consistent with this, Yap1-mGL expression first becomes detectable at the late blastula stage (St.11) (Iwamatsu, 2004), before the onset of gastrulation, appearing as widespread and gradually increasing signal distributed in both the cytoplasm and nucleus of the blastomeres (Figure 1A; Supplementary Movie 1). As epiboly commences (St. 13), Yap1-mGL localisation is observed in the nuclei of the enveloping layer (EVL) and the deep cells, with no detectable fluorescence in the multi-nucleated yolk-syncytial layer (YSL) (Figure 1A; Supplementary Movie 1 and 2). Interestingly, the cells within the domain participating in the rhythmic contractions accompanying gastrulation in medaka display nuclear localisation of Yap1-mGL (Supplementary Movie 1, 2).

**Figure 1.**
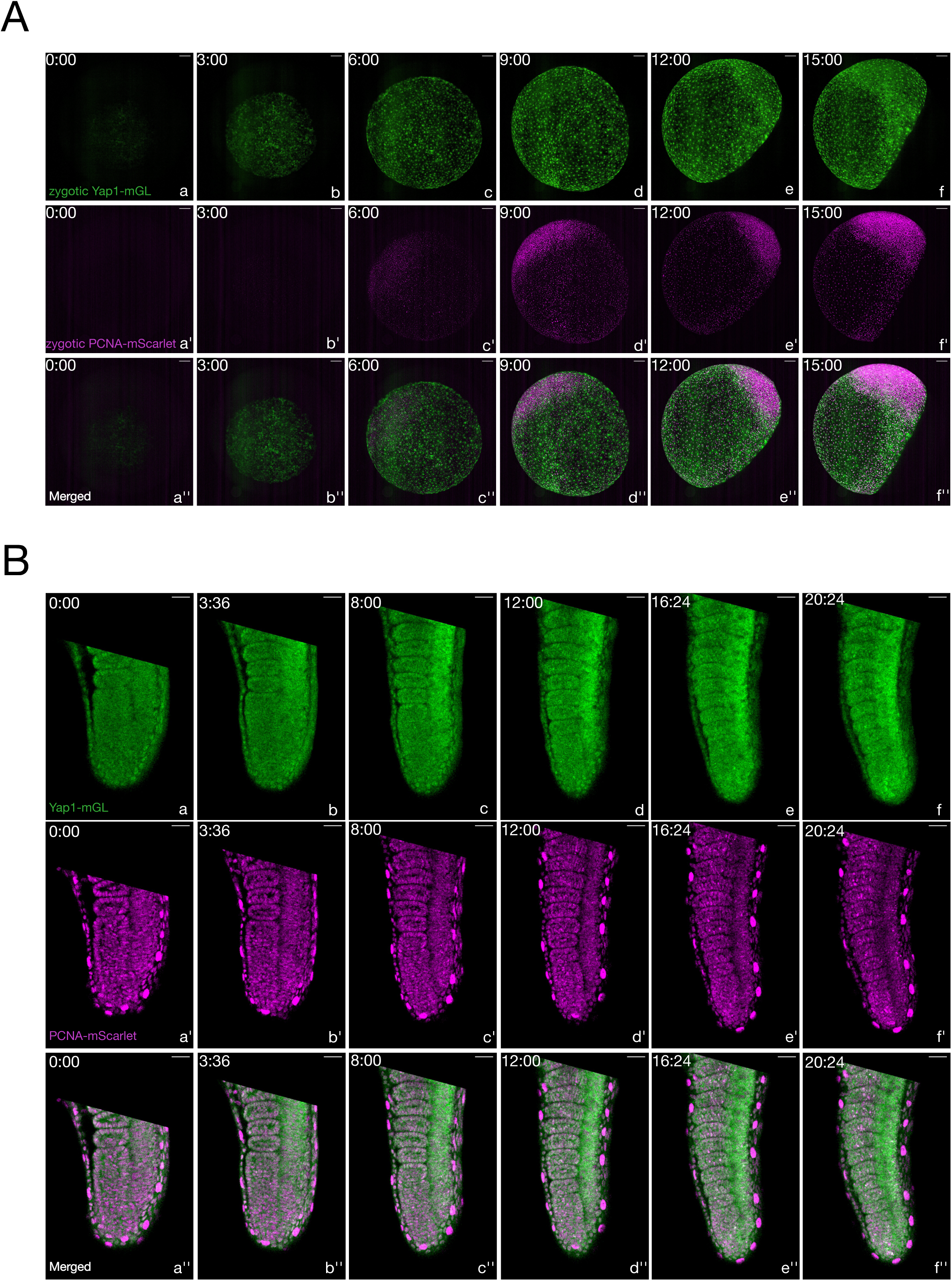
4D imaging of endogenous Yap1-mGL dynamics during gastrulation and somitogenesis. (A) Yap1-mGL (green) fluorescence signal is first detected at St.11 (a) by the late blastula stage; blastomeres display increasing Yap1-mGL expression in an initially salt-and-pepper-like fashion (b) As epiboly and gastrulation commence (c-f), increasing nuclear localisation of Yap1-mGL signal is observed in the enveloping layer (EVL) cells and later-on, in the ventral portions of the embryo participating in the medaka rhythmic contraction waves. Zygotic PCNA-mScarlet expression is detectable after the Yap1-mGL signal is visible (a’-b’). During epiboly and gastrulation stages (c’-f’), widespread nuclear PCNA-mScarlet signal is evident throughout the blastoderm cells except for the YSL and EVL. The differences in expression of both endogenous markers can be better observed in the merged view (a’’-f’’) n = 4 scale bar = 100µm. Time in hh:mm. (B) 4D imaging of tail explants of medaka embryos at the 20-somite stage. Zygotic Yap1-mGL (green) shows widespread expression in the tail (a-f), localisation is predominantly nuclear in forming somites and the enveloping epidermis and primarily cytoplasmic in the presomitic mesoderm (PSM). Maternal PCNA-mScarlet shows widespread nuclear localisation (a’-f’) in forming somites, epidermis, the PSM and the neural tube. The overlap of both endogenous reporters is shown in (a’’-f’’). n=3 Scale bar= 25 µm. Time in hh:mm.

We next assessed Yap1-mGL spatio-temporal expression and subcellular localisation during the process of somitogenesis (Miao and Pourquié, 2024). In mouse PSM cells cultured *in vitro*, high Yap activity suppresses segmentation clock oscillations, whereas lowering Yap signalling switches cells into an oscillatory state, identifying Yap-dependent mechanical cues as potential input to the segmentation clock (Hubaud et al., 2017). By performing 4D time-lapse-imaging on heterozygous PCNA-mScarlet/Yap1-mGL tail explants at the 20 somite stage we report that endogenous Yap1-mGL protein is widely expressed in the developing tail, with expression both in the somites and the unsegmented PSM (Figure 1B; Supplementary Movies 3 and 4). The localisation of the Yap1-mGL signal was qualitatively different, with somites showing a primarily nuclear signal and the unsegmented PSM showing a largely cytoplasmic signal. This reinforces the need to resolve Yap1 sub-cellular localisation even within the same tissue (Figure 1B; Supplementary Movies 3 and 4). Taken together, the Yap1-mGL KI line enables dynamic *in vivo* assessment of Yap1 expression and subcellular localisation during embryonic development in a vertebrate model.

### Quantitative analysis of nuclear/cytoplasmic ratio of Yap1 protein *in vivo*

To enable a more quantitative *in vivo* analysis of Yap1 protein subcellular localisation we crossed the endogenous PCNA-mScarlet reporter to Yap1-mGL and imaged double-positive embryos in 3D with high spatial resolution (Figure 2A-B). We segmented nuclei using the PCNA-mScarlet signal and computed the per cell nuclear/cytoplasmic ratio of Yap1-mGL (Supplementary Movie 5). The per cell and local heatmap of Yap1 nuclear/cytoplasmic (N/C) ratio allow us to quantitatively assess localisation across the intact tail tissue (Figure 2C; Supplementary Movie 5 and 6) (details in Materials and Methods). Our results show that the epidermis, notochord progenitors and somites exhibit a high Yap1-mGL N/C ratio, indicative of active Yap1 signalling, whereas the unsegmented PSM and neural tube show a low N/C ratio consistent with inactive Yap signalling (Figure 2C; Supplementary Movie 5 and 6). The high N/C Yap1 ratio in forming somites and low ratio in undifferentiated posterior PSM is consistent with previous work showing that YAP localises to the nucleus in cells on stiff substrates or under high cytoskeletal tension and to the cytoplasm on soft substrates (Aragona et al., 2013; Dupont et al., 2011). In zebrafish, a fibronectin matrix is assembled along nascent somite boundaries as cells epithelialise (Jülich et al., 2009; Koshida et al., 2005) and adhesion to fibronectin has been shown to promote YAP nuclear localisation (Kim and Gumbiner, 2015). Additionally, our live reporter data is in agreement with earlier antibody-based reports of nuclear Yap1 in the epidermis and notochord of zebrafish and medaka embryos (Kimelman et al., 2017). Taken together, the Yap1-mGL knock-in line in combination with endogenously tagged PCNA-mScarlet enables an *in vivo* quantitative assessment of the nuclear/cytoplasmic ratio of Yap1 protein (as a proxy for signalling activity) across the intact tissue.

**Figure 2.**
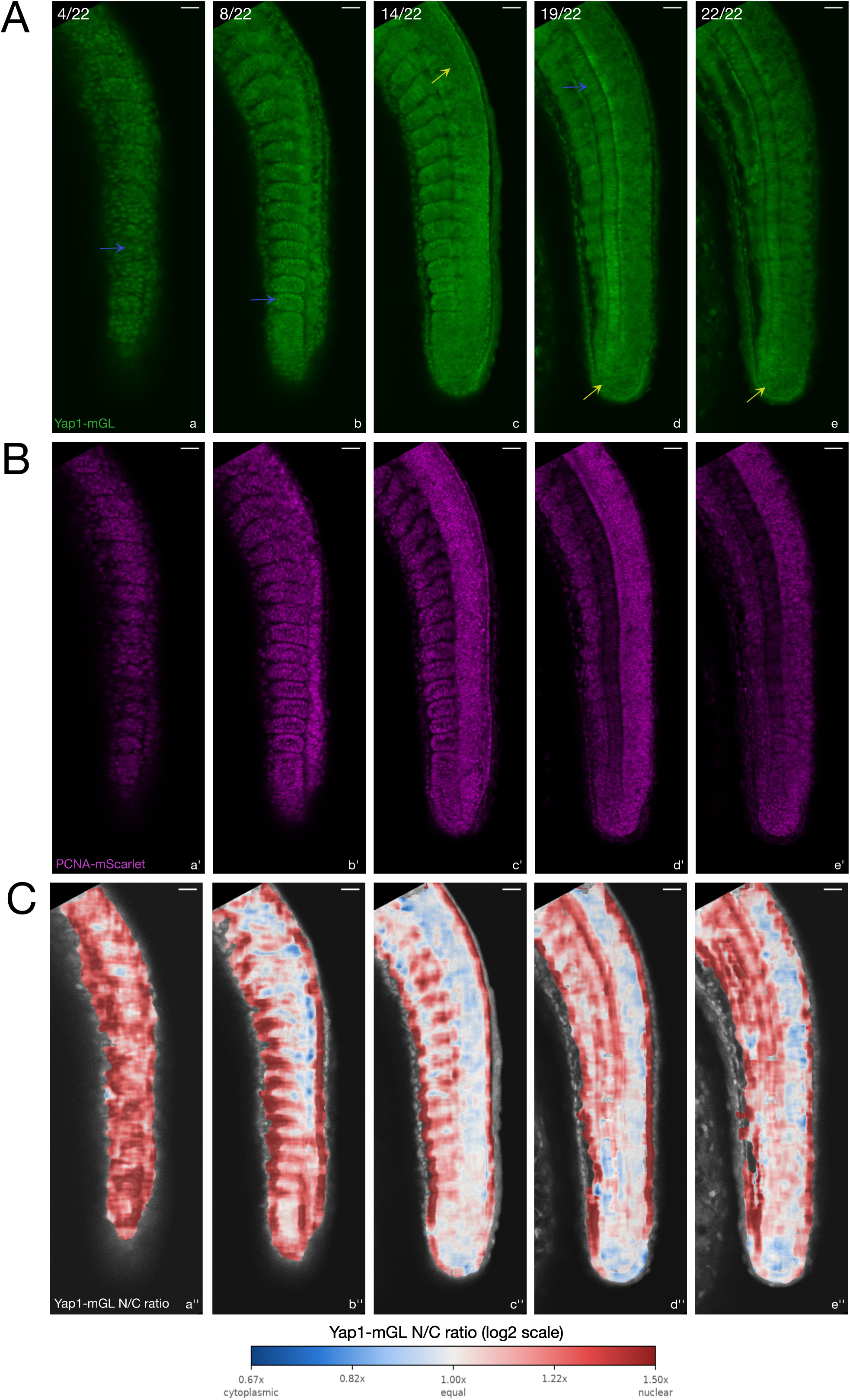
Quantitative analysis of nuclear/cytoplasmic localisation of Yap1 protein *in vivo*. (A) Z-stack through a 20-somite stage embryo. Yap1-mGL (green) shows a predominantly nuclear localisation in the epidermis (blue arrow in a), somites (blue arrow in b) and unipotent progenitors in the notochord (blue arrow in d) and primarily cytoplasmic signal in the neural tube (yellow arrow in c) and posterior PSM (yellow arrows in d-e) (B) PCNA-mScarlet (magenta) shows nuclear localisation across all tissues including epidermis, somites, PSM, neural tube and notochord (a’-e’). (C) Quantitative local heatmap of nuclear/cytoplasmic (N/C) ratio of Yap1-mGL through the tissue confirms a high N/C ratio in the epidermis, somites and unipotent progenitors in the notochord and a low N/C ratio in the posterior PSM and the neural tube, N/C intensity ratio is computed on a log2 scale (blue = cytoplasm enriched, grey = 1, red= nucleus enriched) (a’’-e’’). n=3. Scale bar= 25µm.

### Spatio-temporal dynamics of Yap1 during spinal cord regeneration

Beyond its myriad roles in embryogenesis, upregulation of Yap has been implicated in orchestrating the regenerative response to injury in a variety of models (Moya and Halder, 2019; Riley et al., 2022). Previous work has shown that adult zebrafish fully regenerate the spinal cord after transection through injury-activated Yap signalling in ependymo-radial glial (ERG) cells (Klatt Shaw et al., 2021; Mokalled et al., 2016; Zhou et al., 2023). Adult medaka have been reported to have a limited regenerative capacity after spinal cord transection compared to zebrafish (Aoki et al., 2025), but it remains unclear whether medaka larvae retain their regenerative potential. To assess whether medaka larvae can efficiently regenerate their spinal cord and whether that process includes upregulation of Yap1, we performed a spinal cord transection assay in the Yap1-mGL background. We used two different methodologies to transect the spinal cord in medaka larvae prior to stage 42: a standard injury model and a localised injury model (details in Materials and Methods) (John et al., 2022) (Supplementary Movies 7 and 8). In both models, Yap1-mGL accumulated locally at the injury site relative to the pre-surgery baseline in the same animal (Figure 3A-B). This upregulation was first apparent at 12 hours post-injury (hpi) and persisted to at least 72 hpi with cells upregulating Yap1-mGL spanning the lesion site (Figure 3 A-B). By 72 hpi the lesion site was substantially reduced and tissue continuity was restored, consistent with the structural repair described in regeneration competent zebrafish (Klatt Shaw et al., 2021; Mokalled et al., 2016). Our data suggests that medaka larvae have the potential for spinal cord regeneration and that this process is accompanied by local Yap1 upregulation at the injury site. Given the recently reported limited regenerative response to spinal cord transection in medaka adults (Aoki et al., 2025), how this capacity is attenuated during ontogeny remains to be tested. Overall, our assay shows the utility of using the Yap1-mGL line as an *in vivo* dynamic indicator for Yap1 spatio-temporal expression levels in regenerative assays.

**Figure 3.**
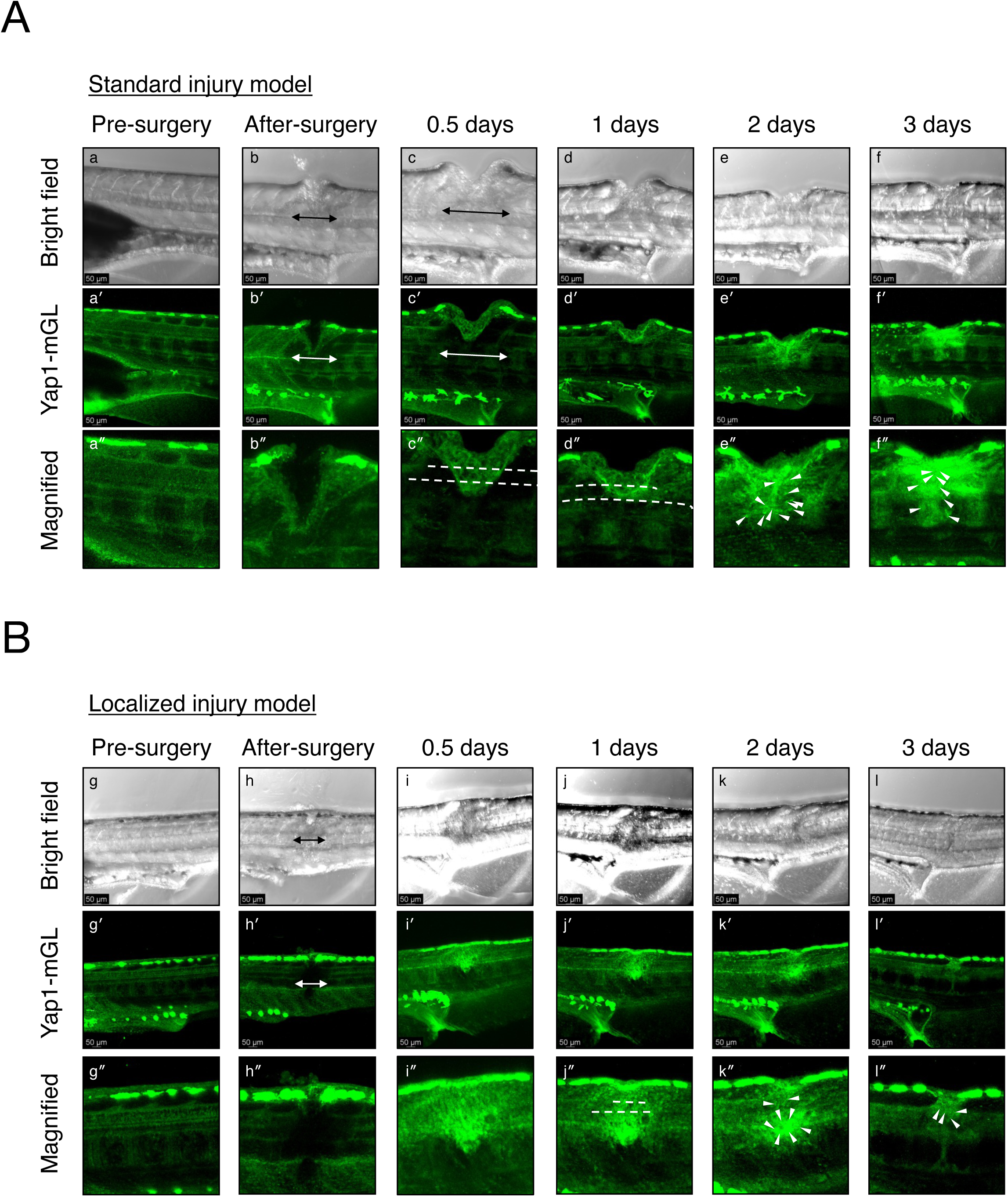
Spatio-temporal dynamics of Yap1-mGL during spinal cord regeneration. Bright-field (top), Yap1-mGL (middle) and magnified Yap1-mGL (bottom) views of the same larva before and after spinal cord transection, in the standard (A) and localised (B) injury model. Larvae were anaesthetised and immobilised under 3% low-melting-point agarose. Double arrows mark the extent of the injured region; dotted lines mark the severed, unconnected neural tube; arrowheads mark accumulating Yap1-mGL-positive cells. Scale bars= 50 µm. n = 7 for each model. (A) Pre-surgery (a–a″): weak Yap1-mGL along the neural tube and intervertebral region. Immediately post-injury (b–b″): the neural tube and surrounding somitic tissue are transected, Yap1-mGL upregulation is visible near the epithelium extending into the injured region by 0.5 days (c–c″), while the neural tube remains severed. At 1 day (d–d″) the wound site is still clearly visible and Yap1-mGL-positive cells appear in the epithelium surrounding the injury and near myotome boundaries. At 2 days (e–e″) neural tube fluorescence is continuous again despite the persisting wound, with Yap1-mGL puncta accumulating around the injured region; this accumulation is maintained at 3 days (f–f″), as is neural tube continuity. (B) Pre-surgery (g–g″): as in A. Immediately post-injury (h–h″): the lesion is largely confined to the neural tube, with limited damage to adjacent tissues such as the somites. At 0.5 days (i–i″) the cord is fully severed but a cluster of Yap1-mGL-positive cells is already present above the injury site, towards the epidermis; the neural tube is not yet reconstituted at 1 day (j–j″). At 2 days (k–k″) neural tube fluorescence is continuous again and numerous Yap1-mGL-positive cells still surround the injury site; at 3 days (l–l″) these cells are still present but reduced in number, and neural tube continuity is maintained.

### Protein degradation using degron nanobodies phenocopies *hirame* mutant

A major experimental goal of current research is to enable the manipulation of protein levels *in vivo,* ideally in a time-resolved manner, to assess protein function (Shi et al., 2025; Yamaguchi et al., 2019). To this end, we combined our Yap1-mGL KI line with the nanobody-based degron system (Yamaguchi et al., 2019) for degradation of endogenous Yap1 *in vivo* (Figure 4A). First, we generated homozygous Yap1-mGL fish that were viable and did not show any obvious phenotypic abnormalities, in contrast to the severe phenotypes reported in the *hirame* (*yap1*) medaka mutant (Porazinski et al., 2015). Next, we injected zGrad mRNA into one-cell stage homozygous Yap1-mGL embryos (Figure 4B). While zGrad has been used to efficiently degrade proteins fused to GFP in zebrafish (Yamaguchi et al., 2019), its utility in medaka and its ability to work with the mGreenLantern fluorescent protein has thus far not been tested. Our data show that zGrad injected Yap1-mGL homozygous embryos exhibit Yap1-mGL degradation, illustrated by the near-complete loss of mGL fluorescence at St. 23 (Figure 4B). By St. 27 we observe a partial return of the mGL signal in the developing body axis, consistent with gradual depletion of the injected zGrad mRNA and hence the transient degradation of Yap1. Importantly, all zGrad injected embryos carrying the Yap1-mGL allele displayed phenotypes that mirrored the *hirame* (*yap1*) loss-of-function mutant (Porazinski et al., 2015) (Figure 4B). Specifically, we could observe a flattening of the body and optic cup, lens misalignment and a *cardia bifida* phenotype (Porazinski et al., 2015) (Figure 4B; Supplementary Movie 9). Control (wild-type) embryos injected with zGrad mRNA did not show any visible phenotype (Figure 4B). This data demonstrates that zGrad mRNA can degrade Yap1-mGL protein *in vivo* and opens up the possibility of targeting other endogenously tagged proteins in medaka (Seleit et al., 2021) using the same methodology. Because the target is the protein rather than the transcript, this approach avoids the transcriptional adaptation that mutant mRNA decay can trigger (El-Brolosy et al., 2019; Rossi et al., 2015), and hence provides an alternative assessment of the functional role of targeted proteins. Lastly, expressing zGrad under tissue-specific or light-induced activation would allow spatially and temporally restricted degradation (Rustarazo-Calvo et al., 2026), a feat that is difficult to achieve using classical genetic mutants.

**Figure 4.**
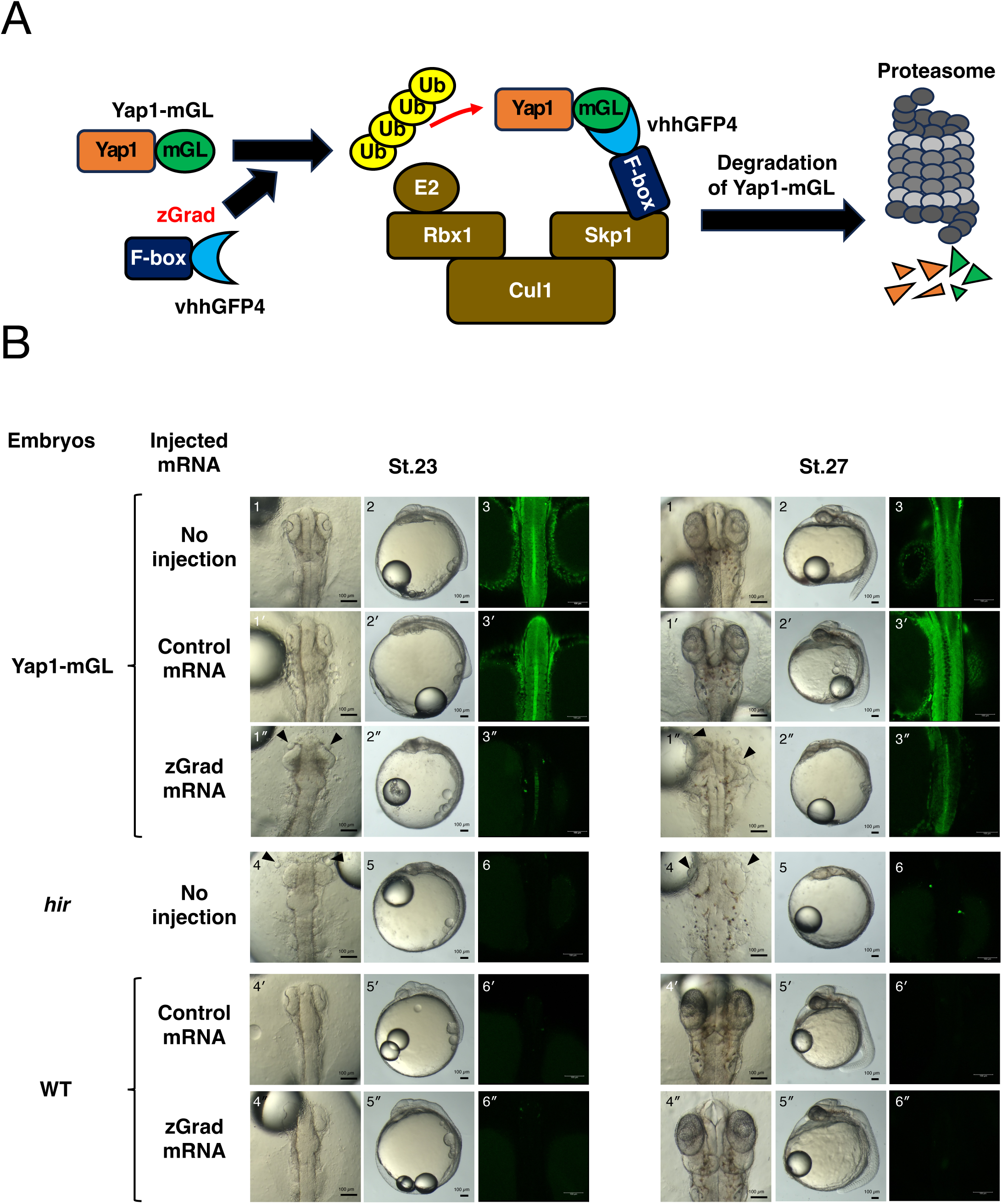
zGrad mRNA injection into Yap1-mGL embryos phenocopies *hirame* (*yap1*) mutant. (A) Schematic of zGrad-mediated Yap1-mGL degradation. An anti-GFP nanobody (VHH-GFP4) with the F-box domain that recruits SCF E3 ubiquitin ligase complex ubiquitinates Yap1-mGL leading to degradation of Yap1 by the proteasome (B) Representative images of medaka Yap1-mGL homozygous embryos without injection (1-3; n=67), with injection of control mRNA (25ng/µl) (1’-3’; n=31), zGrad mRNA (25ng/µl) (1’’-3’’; n=26), *hirame* embryos (4-6; n=16), and WT embryos with injection of control mRNA (25ng/µl) (4’-6’; n=15), zGrad mRNA (25ng/µl) (4’’-6’’; n=31) at St.23 (Left) and St.27 (Right). Scale bars: 100 µm. (1-1’’, 4-4’’) Bright-field images of head region of embryos. Dorsal view, anterior upwards. Arrowheads, mislocated lenses. (2-2’’, 5-5’’) Bright-field images of whole embryos. Lateral view, anterior to the left. (3-3’’, 6-6’’) Fluorescent images of trunk region of embryos. Dorsal view, anterior upwards. The injection of zGrad mRNA markedly reduces Yap1-mGL fluorescence at St.23 with a partial recovery of signal evident at St.27 (see 3, 3’ vs. 3’’). zGrad-mediated Yap1-mGL degradation recapitulates genetic loss-of-function phenotypes of Yap1, such as lens dislocation (1’’ and 4), body flattening (2’’ and 5), and *cardia bifida* (also shown in Supplementary Movie 9).

In this work, we generate an endogenously tagged Yap1 line in a teleost model. We use this line to resolve its spatio-temporal expression and its subcellular localisation, providing new quantitative information on where and when nuclear/cytoplasmic ratios are changing during vertebrate development. We show that the Yap1-mGL protein is detectable from the blastula stage onwards, and exhibits tissue-specific expression and localisation patterns during gastrulation and somitogenesis. Beyond embryonic stages, we find that medaka larvae mount a spatio-temporally confined increase in Yap1 levels in response to spinal-cord transection and that they recover tissue continuity after injury, revealing a larval regenerative competence that appears to be lost in adults. Finally, by coupling the reporter to the zGrad degron system we achieve near complete depletion of endogenously tagged Yap1-mGL that phenocopies the *hirame* mutant. This approach provides a general strategy for loss-of-function analysis in knock-in reporter lines that complements classical genetic mutants. In the future, we believe that combining this methodology with tissue and stage-specific (in addition to optogenetic) control will allow for the dynamic interrogation of the context-dependent roles of key signalling proteins in development and regeneration.

## Acknowledgements

We would like to thank members of the Aulehla lab for feedback and support throughout this project. The European Molecular Biology Laboratory (EMBL-Heidelberg) Genecore is acknowledged for support in whole genome sequencing (WGS) data acquisition and analysis. In particular, we would like to thank Vladimir Benes and Tobias Rausch for computational work on the WGS data and Mireia Osuna Lopez for help in library preparation for WGS. In addition, we would like to thank all animal-care takers at EMBL (LAR) Heidelberg for excellent support. We would also like to thank Furutani-Seiki lab members, particularly William Dean from University of Bath, for his support for the regeneration experiments. Generative AI tools (Claude Fable 5, Anthropic) were used for help with code generation and review and for grammatical and language review, the authors take full responsibility for all scientific content.

## Funding

This work was supported by

The European Molecular Biology Laboratory (EMBL)

The European Research Council ERC consolidator grant agreement n.866537 to A.A.

The EMBL interdisciplinary Postdoc (EIPOD4) Marie Sklodowska-Curie n.847543) to A.S The Grant from the Japanese Ministry of Education, Culture, Sports, Science, and Technology, 25H01789, 24K02226 to M. F-S, 25K10200 to Y.A.

## Author contributions

A.S: Conceptualisation, Methodology, Validation, Formal analysis, Investigation, Data Curation, Writing - original draft, Visualisation.

Y.A: Validation, Formal analysis, Investigation, Data Curation, Writing

S.K.: Methodology, Investigation, Formal analysis, Data Curation, Writing

A.O.: Methodology, Formal Analysis, and Supervision

C.O.: Methodology, Formal Analysis, Investigation

A.Og.: Validation, Formal analysis, Investigation, Data Curation

L.L: Methodology, Investigation, Data Curation

M.F.S.: Conceptualisation, Methodology, Resources, Writing – original draft preparation, Supervision, Project administration, Funding acquisition.

A.A: Conceptualisation, Methodology, Resources, Writing – original draft preparation, Supervision, Project administration, Funding acquisition.

## Data and materials availability

All data are available within the article, supplementary files and source data files. All materials used in this study are available by request from the corresponding author.

## Materials and Methods

### Husbandry and animal ethics

Medaka *Oryzias latipes* (Cab), the endogenous PCNA-mScarlet (Seleit et al., 2021) and Yap1-mGL reporter lines were maintained as closed stocks in a constant recirculating system at 27-28°C, with a 14hr light / 10hr dark cycle in the EMBL Laboratory Animal Resources (LAR) fish facility. Animal experiments were performed after project approval by the EMBL Institutional Animal Care and Use Committee (IACUC), IACUC project code is 20/001_HD_AA. All animal procedures were done with approval by the Committee for the Ethics of Animal Experiments at the Yamaguchi University School of Medicine, Japan. Experiments were carried out according to relevant guidelines and regulations.

### CRISPR/Cas9 and zGrad mRNA injection

For CRISPR/Cas9 injections in *O.latipes* Cab WT background; we used a synthetic gRNA from Sigma-Aldrich (spyCas9 sgRNA, 3 nmol, HPLC purification, no modification) *GCUUCCUCACAUGGUUAUAG* targeting the C-terminus of the medaka *yap1* locus that we located *in silico* using CCTop (Stemmer et al., 2015). PCR repair donor fragments were designed and prepared as described previously (Seleit et al., 2021). mGreenLantern was amplified from (Plasmid #161912, Addgene) using 33 bp homology arms from the endogenous *yap1* locus on both ends. PCR amplifications were performed using Phusion or Q5 high fidelity DNA polymerase (NEB Q5 Master Mix # M0492L). MinElute PCR Purification Kit (Qiagen #28004) was used for PCR purification. Primers were ordered from Sigma-Aldrich (25 nmol scale, desalted) and contained a Biotin moiety on the 5′ ends. The injection mix consisted of the sgRNA (20 ng/µl), the Cas9 protein from IDT (Alt-R™ S.p. Cas9 V3, glycerol-free, 100µg #10007806) at a final concentration of 250 ng/µl and the donor template (8-10 ng/µl). For injections, male and female medaka were added to the same tank and fertilised eggs were collected. The mix was injected in one-cell staged medaka embryos, and embryos were raised at 28°C in 1xERM (Seleit et al., 2021). Founders were identified by fluorescence screening and confirmed by whole genome sequencing (WGS). zGrad mRNA was synthesised from NotI-linearised pCS2(+)-zGrad (Addgene plasmid #119716) using the mMESSAGE mMACHINE SP6 Transcription Kit (Thermo Fisher Scientific), according to the manufacturer’s instructions. zGrad mRNA was diluted to 25 ng/µL and injected into one-cell-stage Yap1-mGL homozygous or wild-type embryos. Phenol red (0.05%; Sigma-Aldrich) was added to the injection solution as a tracer.

### Whole genome sequencing of Yap1-mGL

10 F2 *yap1-mGL* heterozygous embryos were snap frozen in liquid nitrogen in 1.5 ml Eppendorf tubes. Genomic DNA was extracted using DNeasy Blood and Tissue Kit (Qiagen #69504). Library preparation, sequencing and analysis were performed as previously reported (Seleit et al., 2021). Briefly, DNA libraries were generated using a Beckman i7 series and the NEBNext Ultra II DNA Library Prep Kit for Illumina (NEB #E7645S). The libraries were then indexed with unique dual barcodes (8 bp long), pooled together and sequenced using an Illumina NextSeq550 instrument with a 150 PE mid-mode in paired-end mode. Sequenced reads were aligned to the *O. latipes* reference genome (*Ensembl* Assembly version ASM223467v1) using BWA version 0.7.17 with default settings (Li and Durbin, 2009). The reference genome was augmented with the *mGL* insert. SAMtools (Li et al., 2009) was used to sort and index the reads after alignment. For Structural Variant discovery, we utilised DELLY v0.8.7 (Rausch et al., 2012). Integrative Genomics Viewer (Thorvaldsdottir et al., 2013) was used to generate plots in Supplementary Figure 1.

### Live-imaging sample preparation, microscopy

Medaka embryos were prepared for live-imaging as previously described (Seleit et al., 2021; Seleit et al., 2024). 1x Tricaine (Sigma-Aldrich #A5040-25G) was used to anaesthetise dechorionated medaka embryos (20 mg/ml – 20x stock solution diluted in 1xERM). Anaesthetised embryos were then mounted using low melting agarose (0.6 to 1%) (Biozyme Plaque Agarose #840101). Imaging was done in 8-well glass-bottomed dishes (Lab-Tek Chambered #1 Borosilicate Coverglass System 155411, T.S). Embryo screening was performed on a Nikon SMZ18 fluorescence stereoscope with GFP/RFP filters. For live-imaging of Yap1-mGL whole embryos Zeiss LSM780 laser-scanning confocal microscopes with a temperature control box and an Argon laser at 488 nm and a 20× plan apo objective and a laser-scanning confocal Leica SP8 (CSU, White Laser) microscope, 20x and 40x objectives were used. For gastrulation-stage live imaging, embryos from a PCNA-mScarlet male crossed to a Yap1-mGL female were collected and enzymatically dechorionated at stage 10 as previously described (Seleit et al., 2022) and then carefully transferred into 0.8% low-melting agarose inside 8-well glass-bottomed dishes (Lab-Tek Chambered #1 Borosilicate Coverglass System 155411, T.S). Embryos were then oriented animal-pole down using an eyelash, and agarose was allowed to solidify. Wells were covered with 1xERM and microscopy was performed on a Nikon Ti-2 CSU W1 SORA Spinning disk confocal microscope using a P-Apo Lambda S 10X/0.45NA objective and a Diode Laser at 488nm and 561nm, maintaining a constant sample temperature of 22°C. Medaka tail explants were imaged in Culture Medium: DMEM/F12 lacking glucose, pyruvate, glutamine, and phenol red (Cell Culture Technologies), supplemented with 2 mM glucose (45% solution in water, Sigma, G8769), 2 mM L-glutamine (Gibco, 25030), 0.2% penicillin-streptomycin (Sigma, P4333), and ∼0.1% w/v bovine serum albumin (BSA) (Millipore, ES009-B). During imaging, samples were incubated at 27°C, with 5% CO₂ (required for long term stability of Culture Medium pH). Prior to imaging, embryos were transferred from ERM medium to CO₂-independent Dissection Medium, which differs from Culture Medium by being supplemented with 10mM (Gibco, 15360-106) and lacking penicillin-streptomycin. Stage 25-27 Medaka embryos were first dechorionated manually with forceps under a light stereoscope, then the tail region (from its tip to the 2-5 last formed somites) was dissected. Imaging was performed in Ibidi µ-Slide 1 Well (Coverslip: #1.5H Glass, cat. no. 82107), inside which tails were placed in silicon micro-inserts (Ibidi, 80406). Samples were imaged with the same Zeiss LSM 780 setup previously mentioned using a 20x objective. For live imaging of zGrad injected embryos, bright-field images were acquired using a Leica MZ APO stereomicroscope, and fluorescence images were acquired using a Leica DMi8 inverted microscope equipped with an SP8 confocal imaging system.

### Spinal cord regeneration assay

Anaesthetised fish (MS-222) were mounted left-side-up on a No. 1 coverslip (24 × 50 mm; Matsunami C024501) in 3.0% low-melting-point agarose containing 0.168 mg/mL MS-222 and 1× penicillin–streptomycin (Fujifilm Wako, 168-23191). The coverslip rested on a thermoelectric cooler (SCM24-hw; Kouizam) on the stereomicroscope stage; cooling intensity was adjusted while monitoring cardiac activity until the agarose solidified, and the preparation was covered with Parafilm to prevent desiccation. Preoperative Z-stacks were acquired on a Leica DMi8/SP8 confocal system with a 20× glycerol-immersion objective. The preparation was then returned to the stereomicroscope and the Parafilm removed; where needed, 0.8% low-melting-point agarose (same tricaine/antibiotic concentrations) was applied over the solidified 3.0% agarose to maintain hydration and a clear operative field. Transmitted and incident illumination was used for visualisation; the cooler, which blocked transmitted light, was temporarily removed during surgery. The spinal cord was transected at the level of the anal opening using microscalpels fashioned from hypodermic needles. Two injury models, both modified from (John et al., 2022), were applied to medaka larvae before stage 42: a standard injury model (based on the incision lesion model) producing a V-shaped defect extending beyond the spinal cord into surrounding tissues (Supplementary Movie 7) and a localised injury model (based on the perforation lesion model) in which surrounding tissue damage was minimised (Supplementary Movie 8). While the former allows for observation of skin healing, the large size of the wound means that the effects of the injury may not be restricted to the spinal cord. On the other hand, the localised wound in the latter injury model enables a more precise recording of the regenerative process specific to spinal cord injury. The principal modification of both models was agarose immobilisation during surgery, which stabilised larvae and enabled accurate comparison of pre- and post-injury confocal images of the same individual. Surgery was performed on a glass slide as above, allowing immediate post-injury confocal imaging; the procedure was also be performed in a glass-bottom dish (IWAKI 3911-035, No. 1 coverslip). After surgery, the preparation was re-covered with Parafilm and a postoperative Z-stack acquired to confirm complete transection; if unconfirmed, additional surgery was performed until transection was verified. The agarose was then carefully removed with a microscalpel and fine forceps under application of sterile balanced salt solution (BSS) to prevent desiccation, fish were then aspirated head-first into a glass pipette and transferred to a six-well plate (CORNING Costar 6-well Cell Culture plate 3516) with 8-9 mL antibiotic-supplemented recovery medium/well at 28°C, maintained individually. For longitudinal imaging, fish were anaesthetised and immobilised but in 1.1% low-melting-point agarose in a glass-bottom dish.

### Image analysis

Open-source ImageJ/Fiji software and/or Python was used for image analysis of microscopy imaging. For the *in vivo* Yap1-mGL nuclear/cytoplasmic ratio calculation, a tissue mask was defined at each z-section by summing the raw Yap1-mGL and PCNA-mScarlet intensities, smoothing (σ = 3 µm) and Otsu thresholding; background, measured outside this mask, was subtracted from both channels, and all subsequent steps were restricted to the mask. For the local heatmap, the PCNA-mScarlet channel was smoothed (sigma = 0.35 um), divided by its local mean (sigma = 10 um) to cancel depth-dependent attenuation, and Otsu-thresholded within the tissue mask to define nuclear pixels. Pixels within 0.85 um of a nuclear pixel formed a guard band excluded from both compartments; the remaining tissue pixels were considered cytoplasmic. The N/C ratio of a section is the mean background-subtracted Yap1-mGL intensity over nuclear pixels divided by that over cytoplasmic pixels. The continuous local N/C map is computed from these pixel-level compartment masks: a 15 um square window is slid across each section and, at every pixel, the mean Yap1a-mGL intensity over nuclear-compartment pixels in the window is divided by the mean over cytoplasmic-compartment pixels. For the per cell N/C ratio, nuclei were segmented in each section from the PCNA-mScarlet channel with StarDist (Weigert et al., 2020) after Gaussian denoising (sigma = 0.35 um) and contrast-limited adaptive histogram equalisation, keeping objects of 4-80 um^2^. For each nucleus, nuclear signal was measured in the nucleus eroded by 0.57 um and cytoplasmic signal in a 2.0 um ring around it from which all nuclei, guard bands (0.85 um away from the nucleus) and non-tissue pixels were excluded; rings were grown simultaneously with collisions resolved at the midline so that neighbouring cells never share pixels. Nuclei whose ring retained less than 10% of the nucleus area were not measured. Per-cell Nuclear/Cytoplasmic ratio was measured from the remaining nuclei, using fluorescence from the background-subtracted Yap1-mGL channel. Both maps use a diverging blue-grey-red scale that is linear in log2 (N/C), centered on N/C = 1, colour-bar ticks are labelled with the corresponding N/C ratio. Figures were produced with *Matplotlib* (Hunter, 2007) and *pandas* (McKinney, 2010); all image processing used *scikit-image* (van der Walt et al., 2014), *NumPy* (Harris et al., 2020) and *SciPy* (Virtanen et al., 2020). For confocal images of the spinal cord, Leica Image File (LIF) data conversion and metadata extraction were performed using a custom Python program based on *liffile and tifffile* (Christoph Gohlke, 2026) and *NumPy* (Harris et al., 2020). *liffile* was used to retrieve LIF metadata, including spatial-coordinate information; *readlif* was used to access individual image stacks and frames; *NumPy* was used for multidimensional array processing; and *tifffile* was used for TIFF export. Three-dimensional TIFF visualisation and orthogonal-section reconstruction were performed using a separate custom Python program based on *tifffile* (Christoph Gohlke, 2026), *NumPy* (Harris et al., 2020), *SciPy* (Virtanen et al., 2020), and *Matplotlib* (Hunter, 2007). Channel intensity scaling and colour compositing were performed using *NumPy*, and image rendering and figure export were performed using *Matplotlib*. Python 3.13.5 was used.

## Supplementary Figure Legends

**Supplementary Figure 1.**
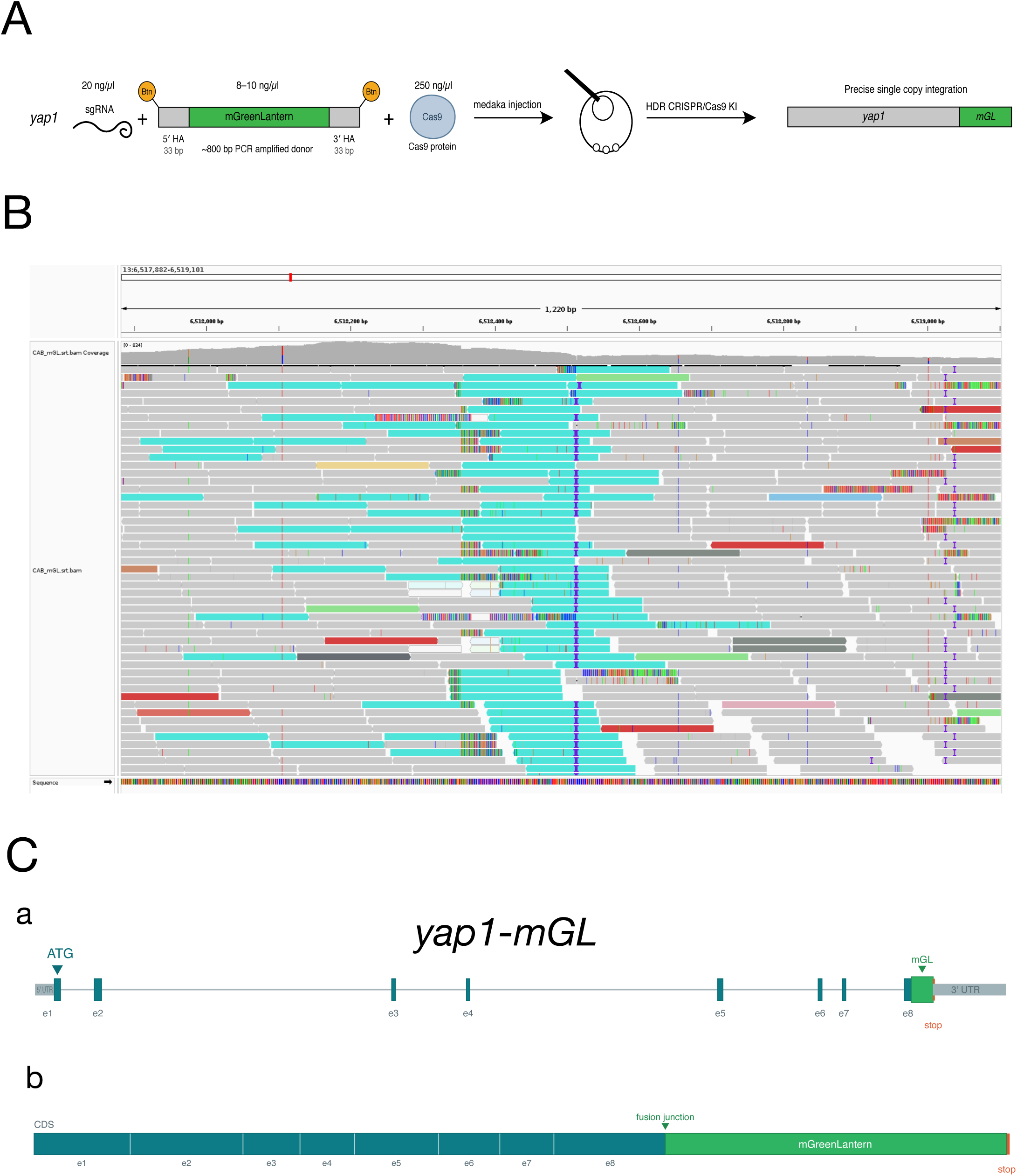
(A) Schematic diagram of the CRISPR/Cas9 knock-in (KI) strategy to tag the endogenous *yap1* locus in medaka. A single-guide RNA (sgRNA) targeting the C-terminus of *yap1* was injected alongside Cas9 protein and the PCR-amplified donor fragment composed of 33bp short homology arms on both ends flanked by the mGreenLantern sequence (with no ATG and no stop codon). Note that the 5′ ends of the PCR donor fragment are biotinylated (Btn). The mix is injected in one-cell staged medaka embryos and the injected fishes were raised and screened for germline transmitted in-frame integrations mediated by homology-directed repair (HDR) (B) mGreenLantern (mGL) integration into the *yap1* locus (Chr13:6,518,353-6,518,362) in *Oryzias latipes*. Paired-end sequenced reads from *yap1-mGL* F2 embryos coloured in grey for concordant mappings and coloured in turquoise for inter-chromosomal paired-ends with one mate mapped to *chr13* and the other mapped to the mGL sequence. The predicted integration site is shown as a vertical dashed line with soft-clipped reads (shown as coloured bases) to the left and right of the integration site. Above the sequenced reads are the basepair-level coverage histogram. The results indicate that *mGL* integrated only at the *yap1* locus genome-wide as we could not detect *mGL* reads anchored anywhere else in the genome. (C) Schematic diagram of the *yap1* genomic locus (a) with exons (e1-e8) in teal, 5′ and 3′ UTRs in grey, mGreenLantern (mGL) in green and the stop codon in orange (b) The coding sequence (CDS) of the Yap1-mGL fusion with the 8 exons in teal, mGreenLantern in green and stop in orange.

## Supplementary Movies Legends

**Supplementary Movie 1** Medaka embryo imaged from late blastula (St. 11) to mid-gastrulation (St. 15). Yap1-mGL (green) fluorescence signal becomes detectable around St.11 in the blastomeres, zygotic PCNA-mScarlet begins to be observed 3 hours later. Yap1-mGL expression is widespread during early gastrulation stages and progressively localises to the nucleus in the EVL and ventral portions of the embryo throughout early stages of epiboly. scale bar= 100µm. Time in hours. n= 4.

**Supplementary Movie 2** Medaka embryo at mid-gastrulation stage (St.15). Animation switching through single channels and a composite view. Yap1-mGL (green) fluorescence signal is widespread at mid-gastrulation (St.15). Zygotic PCNA-mScarlet (magenta) nuclear localisation shows an accumulation of cells at the dorsal presumptive body axis region (top right), ventral (bottom left) cell populations display prominent nuclear localisation of Yap1, with larger EVL nuclei being PCNA-mScarlet negative and ventral ectoderm cells being PCNA-mScarlet positive. n= 4 scale bar= 500µm.

**Supplementary Movie 3** Tail explants of 20 somite stage medaka embryos. Maternal PCNA-mScarlet (magenta) nuclear localisation is visible in the developing somites, neural tube, epidermis and PSM. zygotic Yap1-mGL (green) localisation is visible as primarily nuclear in the developing somites and the epidermis and as predominantly cytoplasmic in the unsegmented PSM. scale bar= 25µm. n=3. Time in hours.

**Supplementary Movie 4** Z-stack through a 10-somite stage embryo. Yap1-mGL (green) shows primarily nuclear localisation in the epidermis, somites and unipotent progenitors of the notochord and a predominantly cytoplasmic signal in the PSM, the notochordal bulb and neural tube. Scale bar= 25µm, n=3.

**Supplementary Movie 5** Quantitative analysis of per cell nuclear/cytoplasmic (N/C) ratio of Yap1-mGL at the 20-somite stage confirms a high N/C ratio in the epidermis, somites and unipotent progenitors in the notochord and a low N/C ratio in the PSM and the neural tube, N/C intensity ratio is computed on a log2 scale (blue = cytoplasm enriched, grey = 1, red= nucleus enriched) n=3 Scale bar= 25µm.

**Supplementary Movie 6** Z-stack through a 20-somite stage embryo. PCNA-mScarlet (magenta) shows nuclear localisation throughout all tissues including epidermis, somites, PSM, neural tube and notochord (left panel). Yap1-mGL (green) shows widespread expression in the tail but distinct localisation of the Yap1 signal, with nuclear localisation in the epidermis, somites and unipotent progenitors in the notochord and cytoplasmic signal in the PSM and the neural tube (middle panel). Quantitative analysis and local heatmap of nuclear/cytoplasmic (N/C) ratio of Yap1-mGL confirms a high N/C ratio in the epidermis, somites and unipotent progenitors in the notochord and a low N/C ratio in the PSM and the neural tube, N/C intensity ratio is computed on a log2 scale (blue = cytoplasm enriched, grey = 1, red= nucleus enriched) n=3 Scale bar= 25µm.

**Supplementary Movie 7** Representative stereomicroscopic recording of the standard spinal cord injury procedure in a medaka larva prior to stage 42. The larva was immobilised in low-melting-point agarose, and an injury was created at the level of the anal opening using a microscalpel fashioned from a 30-gauge hypodermic needle. The microscalpel was inserted laterally, producing a V-shaped lesion that extended through the spinal cord into the surrounding tissues. This model was modified from the incision lesion described by (John et al., 2022) n= 7.

**Supplementary Movie 8** Representative stereomicroscopic recording of the localised spinal cord injury procedure in a medaka larva prior to stage 42. The larva was immobilised in low-melting-point agarose, and the spinal cord was transected at the level of the anal opening using a microscalpel fashioned from a thin insulin needle. The microscalpel was inserted laterally. In contrast to the standard injury model, injury to the surrounding tissues was minimised, resulting in a spatially localised spinal cord lesion. This model was modified from the perforation lesion described by (John et al., 2022) n= 7.

**Supplementary Movie 9** zGrad mRNA injection in Yap1-mGL homozygous embryos phenocopies the *hirame* mutant (including the *cardia bifida* phenotype). Left panel: Yap1-mGL embryos without injection at St.27. The beating heart is located at the center of the embryo. Middle panel Yap1-mGL homozygous embryos at St.27 injected with zGrad mRNA at the 1-cell stage (25ng/µl). Two spatially separated hearts are present, both exhibiting rhythmic contractions phenocopying the *hirame* mutants. Right panel *hirame* embryos at St.27 showing the typical *cardia bifida* phenotype. scale bar= 200µm.

